# ‘Caroten-omics’ of pigments synthesised by *Gordonia rubropertincta* through High Resolution Mass Spectrometry and *in-silico* programming approach

**DOI:** 10.1101/2025.02.18.638947

**Authors:** Gayathri Vemparala, Gangagni Rao Anupoju, Thenkrishnan Kumaraguru

## Abstract

Carotenoids are tetraterpenoid pigments synthesised by microorganisms, plants and present in some animals. They are metabolites of importance in research and industry. The present study aimed to analyse the High Resolution Mass Spectrometry (HRMS) experimental data through R programming and identify all the carotenoids that *Gordonia rubropertincta* produces. The growth rate of the bacterium was 0.24 ± 0.132 h^-^1, produced about 140 ± 20 mg carotenoids/g wet cell weight (WCW) on extraction with methanol and the extract was subjected to HRMS. The number of carotenoids the bacterium synthesizes was identified to be 93 and the concentration of few carotenoids at particular time was pinpointed using R. The distinctive features of the bacterium like high number of carotenoids synthesis through various pathways, their fluctuation with respect to time, storage of carotenoids in cell cytoplasm, and its auto-fluorescent property were also discovered.

## 1.0 INTRODUCTION

Nutraceuticals are potential solutions to challenges such as malnutrition (Katata-Seru et al., 2019), disease prevention (Daliu et al., 2019) and an alternate to chemical food additives and enhancers (Munekata et al., 2020). Molecules like amino acids, vitamins, fatty acids, along with metabolites like carotenoids, antioxidants, polyphenols, fall under the nutraceutical umbrella (GNK et al., 2018; Padmavathi, 2018). Carotenoids are a diverse group of pigments that are popular for their therapeutic properties (Cvetković et al., 2022) and thus are important for production at industrial scale. Carotenoid pigments are produced by wide range of organisms starting from bacteria through plants and also found in animals (De Miguel et al., 2001).

Carotenoids are red-yellow coloured, secondary metabolites made up of isoprenoid units (López et al., 2021). They are tetraterpenoids that contain eight isoprene units (Kumari et al., 2020; López et al., 2021). Most carotenoids are hydrocarbons and are highly non-polar compounds while some carotenoids contain a few oxygen atoms that contribute to their partial polarity (Giuffrida et al., 2018). More than 1200 varieties of carotenoid pigments have been identified to date from various sources and documented (Nupur et al., 2016; Yabuzaki, 2017). Carotenoids are microbial and plant secondary metabolites of high importance in various industries and research purposes. They are well-known for their applications as nutrient supplements, precursors, anti-microbial, as well as cell and DNA protecting agents (Kłodawska et al., 2019). Some carotenoids such as, staphyloxanthin, are known to confer virulence to the host organism producing it. Therefore it can be used as a target for such pathogenic treatments (Ye et al., 2022). According to Business Communications Company (BCC) Research, the current market value of carotenoids is estimated to be $2 billion in 2022 which could rise up to $2.7 billion by 2027 (Bcc and Staff, 2022).

Among the different sources of carotenoids, microorganisms such as bacteria (including Actinobacteria) and microalgae are the best sources for industrial scale production when compared to plants and animals (Loh et al., 2020). Microbes grow quickly in large numbers, are easy to handle at both laboratory and commercial scales, and, simple to study and rewire genetically for higher yield. Microorganisms belonging to genera, *Dunaliella, Haematococcus, Chlorella, Paracoccus, Dietzia, Rhodotorula,* and *Brevundimonas,* are known for their carotenoid producing capacity (Lee et al., 2022; Mussagy et al., 2021; Ram et al., 2020; Zhao et al., 2023). Actinobacteria contain bacterial species that are mostly pigmented and thus appear coloured. Genera such as *Micrococcus, Arthrobacter, Curtobacterium, Corynebacterium,* and *Rhodococcus* are actinobacteria that are known to produce carotenoids (Sandmann, 2021).

*Gordonia* is an actinobacterial genera that were reported to synthesise new carotenoid types that were not known previously (Sowani et al., 2018). The *Gordonia* species like *G. alkanivornas* (Silva et al., 2016)*, G. ajoucoccus* (Kim et al., 2014)*, G. tearrae* (Elfalah et al., 2013), and *G. jacobaea* (De Miguel et al., 2001), are known for their ability to synthesize carotenoids. *G. rubropertincta* is another species that was not studied with respect to the carotenoid pigments it synthesises. Moreover, to study carotenoids synthesised by any bacterium, elaborate extraction procedures and analysis such as High Performance Liquid Chromatography (HPLC) that require standardizations with controls is necessary which might make the process expensive and limited to certain carotenoid metabolites. Therefore, the intention behind current work is to identify the carotenoids that *G. rubropertincta*, produces by combining the methods of untargeted metabolite analysis through High Resolution Mass Spectrometry (HRMS) and the data analysis using R programming to quickly identify all carotenoids in the bacterium.

Initially, the growth pattern of the bacterium was studied which gave us a brief insight into how the pigment synthesis might proceed with respect to the incubation time as microbes synthesize carotenoids during the stationary phase of growth.. Then a solvent extraction method was applied to extract the pigments from the bacterial cell. Then, extracted pigments were identified by HRMS which revealed various types of carotenoids pigments the bacterium synthesises. The data obtained through HRMS were sorted and interpreted. Finally, the morphology of the microbe was studied initially by Scanning Electron Microscopy (SEM) followed by fluorescence and confocal microscopy to locate the carotenoids in the cell. Transmission Electron Microscopiy (TEM) technique was employed to confirm the location of the pigment in the cells which was expected to aid in the carotenoid extraction process.

## 2.0 MATERIALS AND METHODS

### 2.1 Bacterium acquisition, maintenance, and morphological studies

The current study was carried out using *G. rubropertincta* collected from Microbial Type Culture Collection and Gene Bank (MTCC), CSIR-Institute of Microbial Technology, Chandigarh, India, (Accession No. 289). The microbial culture was maintained in the Nutrient Broth medium (Himedia). For all the experiments of the present study, 1000 mL of bacterial culture was grown at 25 °C for 48 h and stored at 2-4 °C.

### 2.2 Bacterial growth curve and growth rate analysis

To perform this experiment, bacterial culture was inoculated in nutrient broth medium, and 1 mL of liquid sample was collected every 2 h up to 48 h and later, samples were collected every 24 h. Based on the results the experiment was terminated after 15 d. The culture was centrifuged at 7000 rpm for 2 min. The supernatant was discarded, and the pellet weight was recorded to calculate growth rate of the microbe. The pellet was then suspended in distilled water and Optical Density (OD) was taken at 600nm (Hach-DR6000 UV-VIS spectrophotometer, Loveland, CO, USA). Graph has been plotted for growth curve and growth rate was calculated using the Equation 1 (Fernández-naveira et al., 2016). This experiment was performed in triplicates and the average of these triplicates were used to plot the graphs.

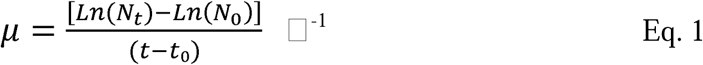

Here, *µ* is the growth rate, *N_t_* is the cell density in g/L at time t (hours), *N*_0_ is the cell density at time 0 h (t_0_).

### 2.3 Extraction and quantification of pigments

The carotenoid pigments were extracted from the bacterial cell at the end of 15 d of incubation through a modified solvent extractions process (Bisht et al., 2020; Poddar et al., 2021). The bacterial culture was pelleted by centrifuging at 7000 rpm for 5 min. The supernatant was removed, and pellet was washed thrice with distilled water. 50 mg of wet pellet was taken in five centrifuge tubes each for four different solvents extraction and a combined solvents extraction. The solvents used for extraction were acetone, methanol, 1-butanol, and ethyl acetate (Sisco Research Laboratories Pvt. Ltd., Mumbai, India). Their combination in equal proportion was also used for extraction. 2 mL of solvent respectively was added to each of the tubes and vortexed for 1-2 min. It was then centrifuged at 7000 rpm for 5 min. The supernatant solvent containing the pigments was collected into separate centrifuge tubes and the procedure was repeated thrice. To quantify the pigments, the solvent was dried by a rotary evaporator (Heidolph Schwabach, Germany) and; remaining pigments were weighed and recorded. After weighing, the pigments were again dissolved in 2 mL respective solvents. All the solvents were mixed and lyophilised. (Labconco-FreeZone Plus 6 lyophiliser). The lyophilised pigments were used for further experiments.

### 2.4 Liquid Chromatography/High Resolution Tandem Mass Spectrometry (LC/HRMS)

The lyophilized pigments were dissolved in methanol and the types of carotenoids present in the sample were analysed by LC/HRMS (Waters-Xevo G2-XS QTof, Milford, MA, USA). This method was developed for the identification of carotenoid pigments and was done by using the data about every carotenoid such as molecular weight and structure available in the Carotenoid Database (López et al., 2021).

### 2.5 Scanning Electron Microscopy (SEM)

The bacterial culture was centrifuged as above to get a pellet and supernatant was discarded. The pellet was washed thrice with distilled water and then lyophilized. The lyophilized bacterial culture was placed on a conductive carbon double-sided adhesive tape (Nisshin EM Co., Ltd., Tokyo, Japan) attached on one side to a metal stub. The stub was left to rest to facilitate the sample sticking to the carbon tape. The stub was placed in the sample chamber of the SEM and observed (Jeol-JSM-7610F Schottky Field Emission Scanning Electron Microscope, Tokyo, Japan).

### 2.6 Fluorescence and Confocal microscopy

The localisation of the pigments synthesised by the bacterium was done using various microscopic techniques. To find out the wavelength range of the pigments produced by the bacteria under study, initially the bacteria were observed under three filters of a fluorescent microscope namely, DAPI, GFP and TRITC. This is a simple procedure designed to fix bacterial cells between a slide and coverslip for making the observation under the microscope easy. The slide was prepared by pelleting the cells by centrifugation, followed by washing the pellet with 1X Phosphate-Bufferd Saline (PBS) for 3 times. The cells were then suspended in 1X PBS and 200 µL was layered onto a sterile 22 mm^2^ cover slip which was placed in a sterile Petri dish. The setup was incubated overnight under 30 °C temperature. Then, the cover slip was washed thrice with 1X PBS and observed under bright-field microscope to confirm the cellular attachment to the cover slip. The cover slip was fixed on to a microscopic glass slide using Eukitt® Quick-hardening mounting medium (Sigma-Aldrich, Burlington, MA, USA). The slide was observed under a fluorescence microscope (Olympus-IX73P1F, Tokyo, Japan) and then under confocal laser scanning microscope (Olympus-FluoView FV10i, Tokyo, Japan).

### 2.7 Transmission Electron Microscopy (TEM)

To confirm the location of the pigments inside the bacterial cell, TEM was employed. The bacterium was grown as described above and pelleted by centrifugation. The pellet was preserved in 2.5% glutaraldehyde (Sigma Aldrich, 0.1M phosphate buffer – pH 7.3) and stored at 4 °C for 24 h and processed for the TEM study as per standard protocol(Lakshman, 2019). The samples were washed in buffer and post fixed in 1% aqueous Osmium tetroxide (Sigma Aldrich) later dehydrated in ascending grades of acetone (Qualigens fine chemicals, Mumbai, India), embedded in Spurr’s resin (SPI supplies, araldite 6005 Embedding kit, USA) and were incubated (Universal incubator-NSW-151) for 24 h at 70 °C for complete polymerization of tissues. The blocks were trimmed manually and semi thin (700-800 nm) sections were made with ultra-microtome (Leica ultra-cut UCT-GAD/E-100, Germany), stained with 1% toludine blue (Qualigens fine chemicals) and observed under a light microscope (Olympus-AX 70) to locate exact area and to remove extra resin, if any for making ultra-thin sections (50nm thickness). Suitable ultra-thin sections were mounted on copper grids (SPI supplies, USA) allowed to air dry overnight and were stained with Urenyl Acetate (UA) and 1% Reynold’s Lead Citrate (LC) as previously reported (Bozzola and Russell, 1998). All grids were dried at room temperature and observed under TEM (Talos F200X TEM) at 120kV.

## 3.0 RESULTS

### 3.1 Bacterial growth curve and growth rate analysis

The bacterial growth curve shows the transition of growth phases of the bacterium under study with respect to time. This information should aid in studying the pigment production as carotenoids are secondary metabolites synthesised by the bacterium during its stationary phase (Narsing Rao et al., 2017). The bacteria, as can be seen in **Fig. 1(a)**, remain in the lag phase for about 4 h after inoculation and then enters the logarithmic/exponential phase. The log phase initiates and rapid exponential growth occurs between 6-12 h of incubation and hence a higher deviation in the OD could be seen. After 12 h, although not exponential, the bacterium remains in log phase for about 3 d as can be seen from **Fig. 1(b)**. The stationary phase then starts and continues to persist for a very long time, i.e., up to 15 d which can be noticed from **Fig. 1(b)**, and then starts descending gradually. Hence, the bacterium had the potential to synthesize pigments for about 15 d after inoculation. Also, the maximal growth rate of the bacterium was found to be 0.24 ± 0.132 h^-1^.

**Fig. 1(a):**
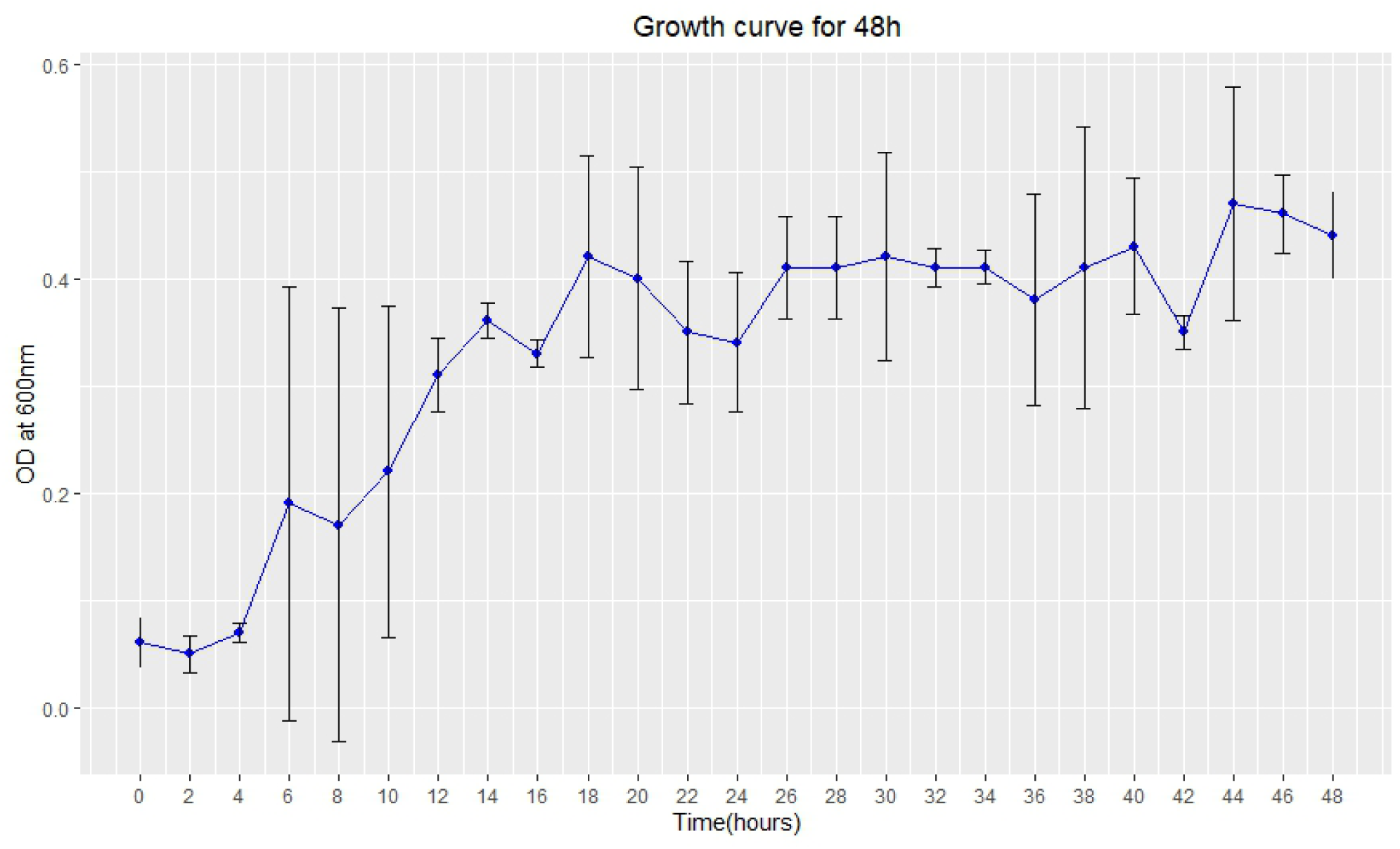
*Gordonia rubropertincta* growth curve for 48h. The dots represent the mean of triplicate experiments and the error bars represent standard deviation from mean.

**Fig. 1(b):**
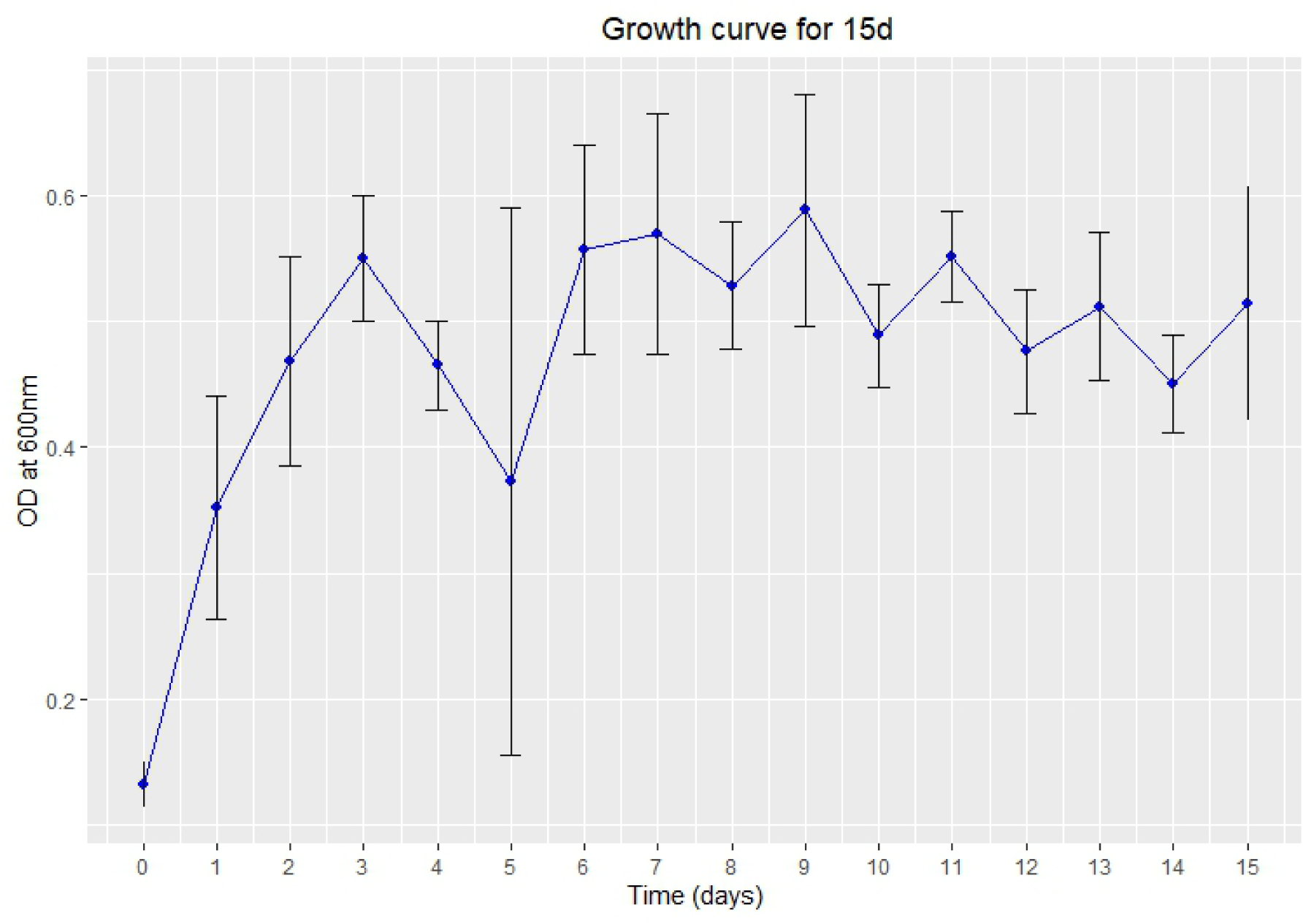
*Gordonia rubropertincta* growth curve for 15d. The dots represent the mean of triplicate experiments and the error bars represent standard deviation from mean.

### 3.2 Extraction and quantification of pigments

The pigment extractions were performed by solvent extraction method at the end of 15 d of incubation. Four solvents were chosen based on their polarity, boiling points, pigment solubility, ease of availability and solvent recovery. A combination of all the solvents in equal proportion was also used. Acetone, methanol, 1-butanol, ethyl acetate, and the combination yielded an average of 100 ± 20mg, 140 ± 20mg, 66.67 ± 16mg, 126.67 ± 12.5mg and 90.67 ± 19mg carotenoids/g wet cell weight (WCW) respectively. Fig. 2 is a graphical portrayal of the pigment yield with various solvents. Of all the solvents, it is evident that methanol could extract the highest amount of pigment compared to the other solvents making it the most suitable solvent for carotenoid pigment extraction from the bacterium, *G. rubropertincta*.

**Fig. 2:**
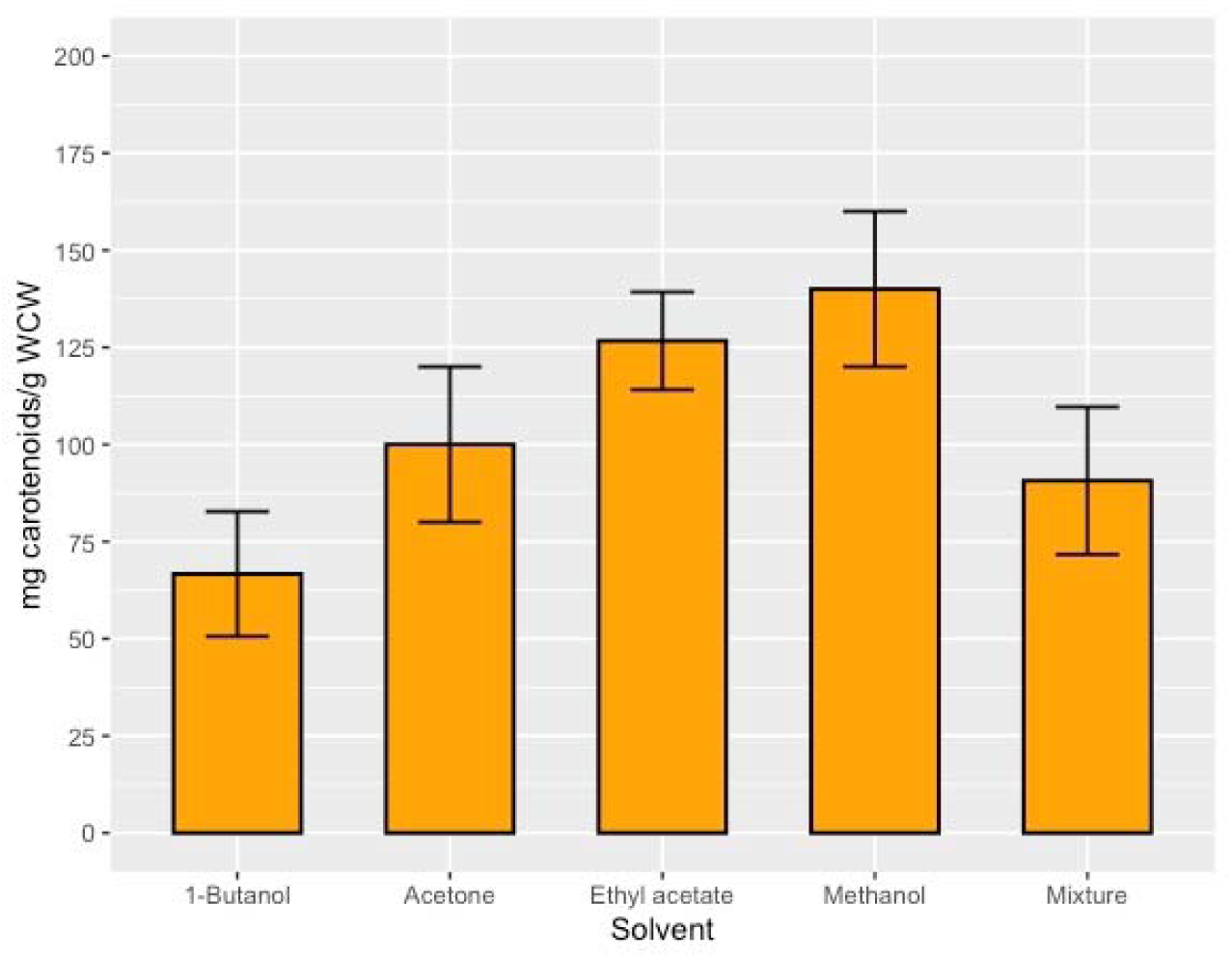
Carotenoids extracted from *G. rubropertincta* using various solvents and their quantification. The bars are plotted using mean of triplicate experiments and the error bars represent standard deviation from mean. The results are statistically significant with a p-value < 0.05.

### 3.3 Liquid Chromatography/High Resolution Tandem Mass Spectrometry (LC/HRMS)

As the types of carotenoid pigments present in the bacterium are unknown, a whole metabolite analysis was performed using the HRMS technique. The analysis revealed that the bacterium synthesises 93 types of carotenoid pigments whose quantity in terms of percentage can be seen in **Fig. 3(a)** and out of them, 16 accounted for highest quantity than others. The obtained results were compared with the KEGG carotenoid biosynthesis pathway database (https://www.genome.jp/kegg/pathway.html) and it was determined that the high number of carotenoids identification through HRMS is due to the fact that the intermediates of the pigments synthesis pathways are also carotenoid type. Therefore, the intermediates and the end products together resulted in a very high number. All the identified pigments and their abundance could be seen in **Figure S1**. To confirm this interpretation, same analysis was performed with pigments extracted at specified time intervals of 3 days up to 15 days to check the progression of carotenoid synthesis. The obtained HRMS data were sorted based on factors like carotenoids present throughout 15 d, carotenoids specific to a particular time interval but not present on other days, the quantities of each carotenoid and their fluctuations throughout 15 d. The detailed information on the carotenoids and their abundance on each day are given in the **Figure S2(a)-(e)** in the **supplementary material**. **Fig. 3(b)** shows only those carotenoids that are present continuously throughout all the 15 days and how their quantities fluctuate with time. The presence of varieties of carotenoids varied in quantity at different times of bacterial incubation shows that this is a simple and effective combination of methods to determine the concentration of any metabolite. Also, the bacterium under study is a store house of wide range of carotenoids that could have multiple applications in research and industry.

**Fig. 3(a):**
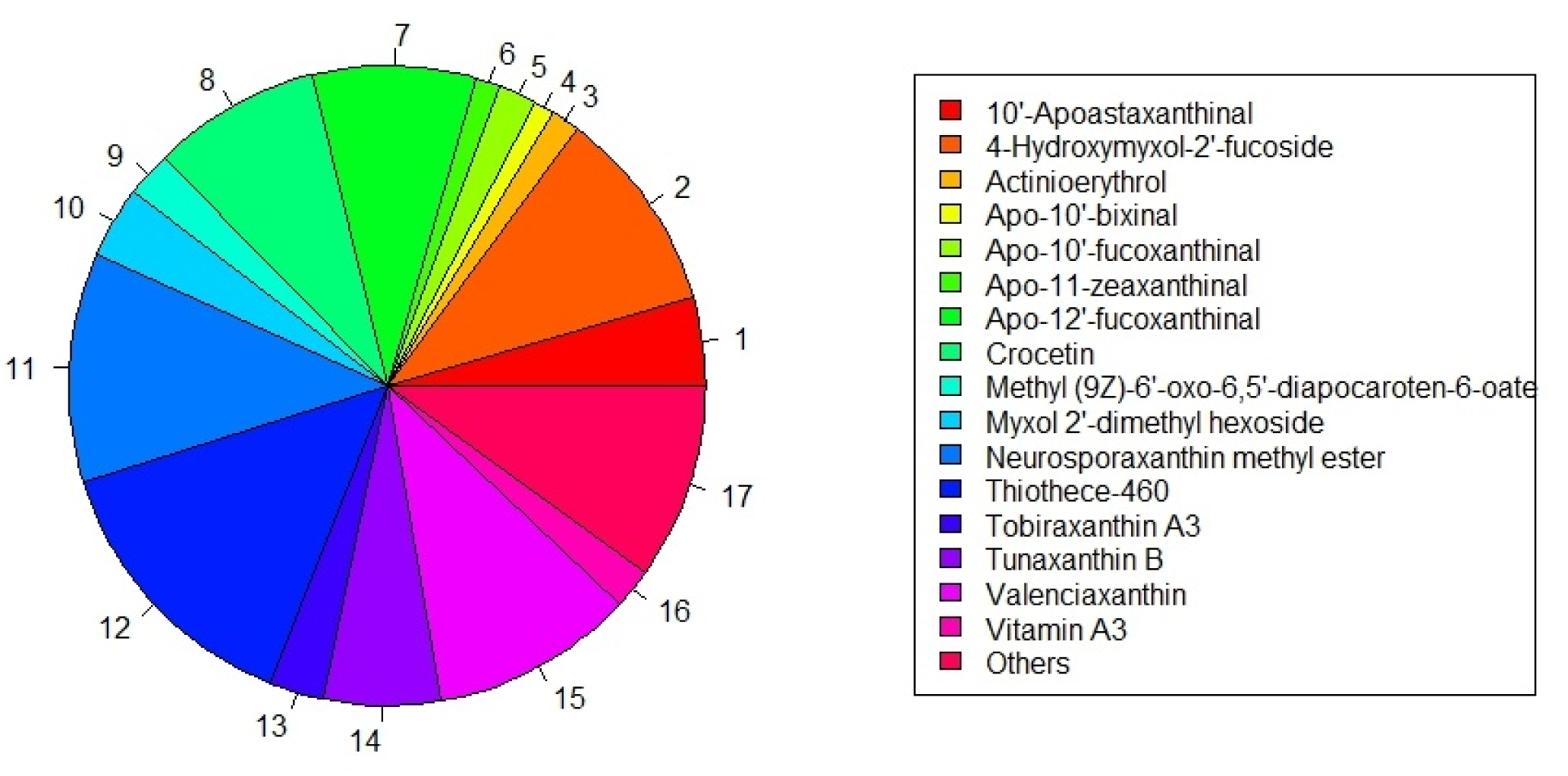
Carotenoid pigments synthesized by *G. rubropertincta* in highest quantity identified by High Resolution Mass Spectrometry

**Fig. 3(b):**
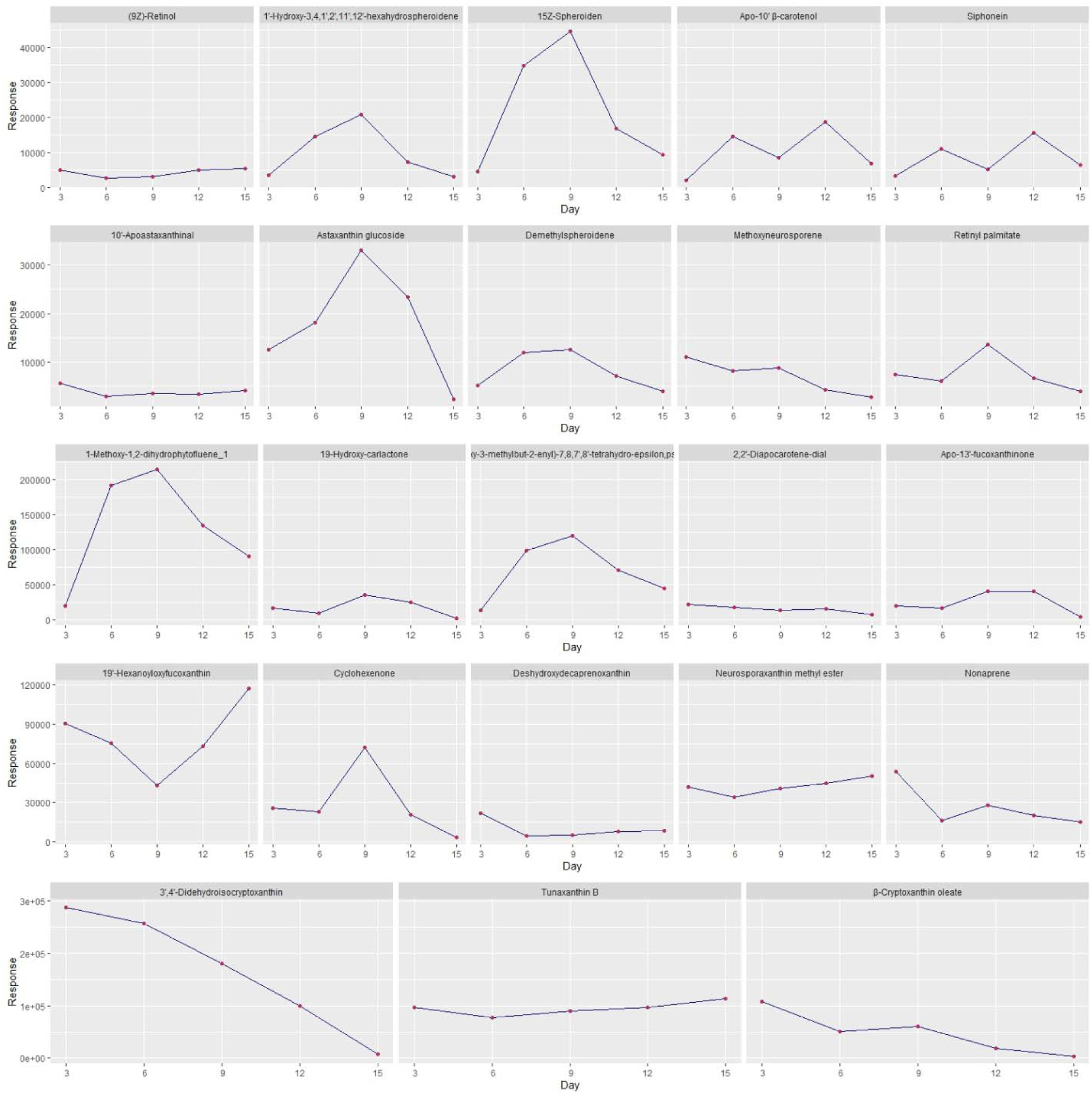
Carotenoid pigments synthesized by *G. rubropertincta* that are present throughout the 15 d and their varying concentrations at regular time intervals identified by High Resolution Mass Spectrometry

### 3.4 Scanning Electron Microscopy (SEM)

**Fig. 4** shows microscopic observation of *G. rubropertincta* which revealed the surface morphology to be long, rod-shaped, bacilli-like structures although cells slightly vary in length. The bacterial cells grow at one or either ends attached to another cell creating a film or a layer in the broth medium. This might be due to the formation of non-septate hyphae. The width and depth differ at each point of the microbe resembling a rather non-uniform bacillus. Some cells show a characteristic structure of one end branching into two and giving it the shape of the alphabet letter Y. Evidently, the microbe shows features peculiar to actively growing *Actinobacteria* (Hopwood, 2007), and further experiments in this study uncover some particular cellular details about *G. rubropertincta*.

**Fig. 4:**
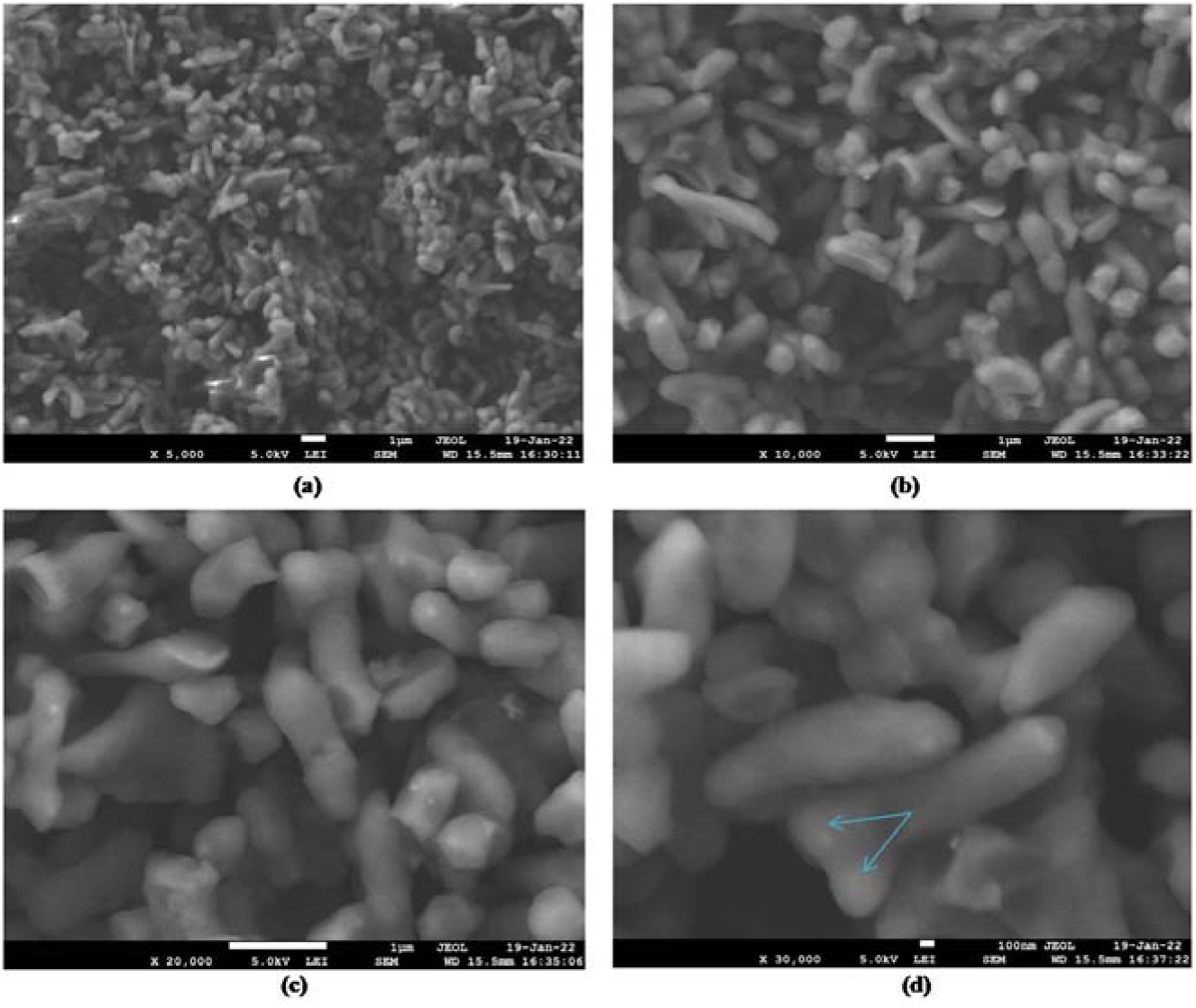
Scanning Electron Microscopy of *Gordonia rubropertincta* (a)-(d) micrographs of increasing magnification. (a) & (b) long, slender and connected bacterial cells (c) cells resemble rods with varying measurements (d) blue arrows indicate cell branching at one end

### 3.5 Fluorescence and Confocal microscopy

The bacterium was observed under three filters, DAPI, GFP and TRITC of a fluorescence microscope to find out the fluorescing capability of the pigments and also to locate them in the cell. The whole cell appeared blue in colour under DAPI filter indicating that the pigments could fluoresce and are carotenoid type. The location of the pigments could not be pinpointed and hence a confocal microscopic technique was employed. **Fig. 5** shows the bacterial cell under confocal microscope with carotenoids fluorescing in blue. Two distinct layers could be seen on the exterior of the cell which probably is the cell wall and plasma membrane where no fluorescence could be seen. As the bacterium under study is a gram positive (Nouioui et al., 2018) one the outermost thick layer of the cell can be attributed as the cell wall with a cell membrane inner to it. The fluorescence emission occurs from the interior of the cell and that indicates the pigment location to be the cytoplasm. Further experiments were carried out to confirm this interpretation.

**Fig. 5:**
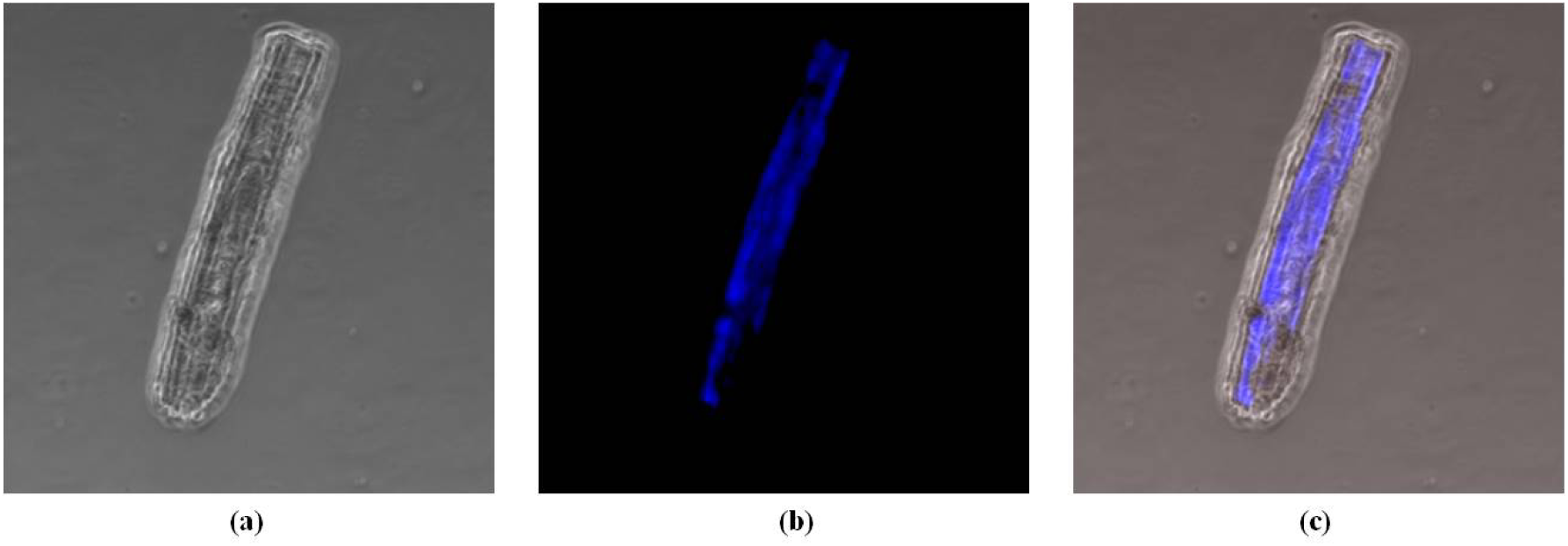
Confocal Microscopy of *Gordonia rubropertincta* (a) Phase contrast image of a single bacterial cell. (b) Fluorescence image showing the pigments (blue fluorescence) inside the cell. (c) Merged image

### 3.6 Transmission Electron Microscopy (TEM)

The TEM image, **Fig. 6**, gave a clear picture of the bacterial structure and the pigment location. The exterior black line in **Fig. 6(b)**, is the stained cell wall and plasma membrane of the bacterium. The genetic material appears as darkly stained body inside the cell as it gets stained by UA (Kabiri et al., 2019) and the whole cell appears dark indicating the presence of carotenoids inside the cell being distributed all over in the cytoplasm. **Fig. 6(a) and (c)** also shows dark cells with two distinct layers outside the cell and the genetic material inside as a black dot. TEM provides evidence that the bacterium under study stores the synthesised carotenoids in the cell cytoplasm. Till date carotenoids were known to be present in bacterial cells bound to their plasma membranes (Hagaggi and Abdul-Raouf, 2023; Kirti et al., 2014). Determining the location of carotenoid storage inside the bacterial cell helps in developing efficient methods to extract the carotenoid pigments from the bacterium.

**Fig. 6:**
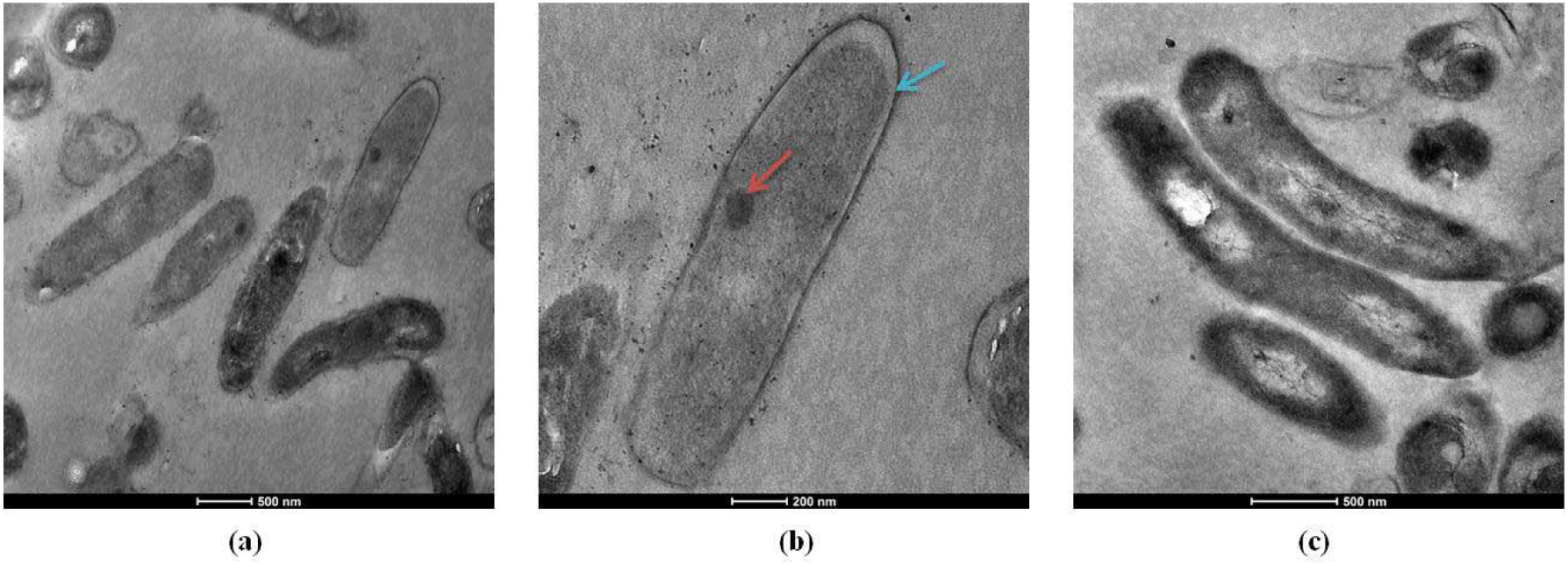
Transmission Electron Microscopy of *Gordonia rubropertincta* (a) & (c) Individual bacterial cells. (b) One bacterial cell with dark cytoplasm. Blue arrow indicates the plasma membrane and red arrow shows the genetic material of the bacterium

## 4.0 DISCUSSION

The carotenoid pigments produced are secondary metabolites and therefore it would be useful to precisely know the stationary phase of the bacterium for extraction of pigments. Of the four growth phases, the bacterium enters the stationary phase between 6-12 h of incubation and remains in it for about 15 d. The growth rate of the bacterium was 0.24 ± 0.132 h^-1^.

Four solvents, acetone, methanol, 1-butanol, ethyl acetate and an equal mixture of all four were used for the extraction process. Carotenoids are C_20_-C_60_ hydrocarbon in nature with some containing a few oxygen atoms. They are highly non-polar molecules with the oxygen contributing to a little polarity to some carotenoids. Hence, solvents of increasing relative polarity (Miller’s Home Solvent polarity table) were chosen for extraction so that carotenoids of various polarities get dissolved in the solvents and are extracted easily. Also, the availability of solvents, their ability to dissolve cell wall and plasma membrane to release cellular contents including carotenoids, solubility of pigments and ease of solvent recovery were other solvent properties taken into consideration. The carotenoids could be easily extracted from the cells and separated from cell debris through centrifugation. Of the solvents, methanol was found to be most suitable for extracting high amount of carotenoids i.e., 140 ± 20 mg carotenoids/g WCW. Thus extracted carotenoids were then lyophilised and used for the identification of types of carotenoids by HRMS method.

A mixture of carotenoids were observed that were 93 in total. Each carotenoid showed a particular response that corresponds to the quantity present in the whole mixture. So, HRMS is both a qualitative and quantitative estimation of whole carotenoids present in the extract. When compared with the KEGG (https://www.genome.jp/kegg/pathway.html) and Carotenoid database (Yabuzaki, 2017) the 93 carotenoids identified were found to be the intermediates and end products of carotenoid synthesis pathways. It was also found that *G. rubropertincta* is a bacterium capable of synthesising varieties of carotenoids some of which are not known till date to be synthesised by bacterium but produced by higher organisms such as algae, fungi and plants. For example, neurosporaxanthin methyl ester (Parra-Rivero et al., 2020) and crocetin (Guo et al., 2022) that have pharmacological applications are generally found in fungi and plants respectively (Yabuzaki, 2017). Synthesis of such important carotenoids by a bacterium could be highly beneficial to produce them at commercial scales.

The HRMS analysis was performed on carotenoids extracted at regular intervals of time for 15 d as the bacterium stays in stationary phase for about 15 days as found through growth curve analysis. Extraction was performed every 3 d up to 15 d. Intermediates and end products were seen in all 5 samples with the types and abundances varying on each day. This information aids in identifying the pathway(s) through which the organism is actively synthesising carotenoids, the concentration of each carotenoid at any point of time, so, targeting and harvesting any particular carotenoid is possible. *G. rubropertincta* synthesises carotenoids through diverse pathways such as Neurosporaxanthin, bacterioruberin, spirilloxanthin, okenone. While the types and concentrations of carotenoids on each are varying, 23 carotenoids were found to be present throughout the 15 d. The time point at which a carotenoid is present at maximum quantity can be identified through this method which helps in deciding the incubation or fermentation time to obtain the maximum amount of product.

SEM revealed the bacteria to be rod-shaped, bacilli-like but cells connected to each other due to the formation of non-septate hyphae. Carotenoid pigments such as, β-carotene, rhodopin, spheroidenone, fucoxanthin, lycopene (Gillbro and Cogdell, 1989; Katoh et al., 1991) are all fluorescent carotenoids whose fluorescence spectra lies in the UV-VIS region (Miller et al., 1935). *G. rubropertincta,* when observed under three different filters i.e, DAPI (4’,6-diamidino-2-phenylindole), TRITC (Tetramethylrhodamine), GFP (Green Fluorescent Protein), of a fluorescence microscope, it fluoresced under the DAPI region showing the carotenoids it synthesises are also auto-fluorescent. Taking advantage of the auto-fluorescence, the bacterium was then observed under a confocal microscope to precisely locate the carotenoids inside the cell as that information can be useful in developing an efficient extraction strategy. When observed using a confocal microscope, a much clearer and sharper image of the bacterium was obtained showing some fine details on the cellular characteristics. The bacterial cell consists of a thick cell wall as it is gram-positive in nature, a lipid bilayer inside the cell wall, and the fluorescence emerges only from the inside of the plasma membrane indicating the location of carotenoids in the cell cytoplasm. It has been previously reported that carotenoids are present in the plasma membrane bound to one of the leaflets of the lipid bilayer or embedded inside the cell membrane with hydrophobic ends tucked inside and exposed hydrophilic ends (Anwar et al., 1977; Chia et al., 2021). Bacterial carotenoids are also found bound to the bacteriochlorophyll pigments present in the plasma membrane (Yurkov et al., 1993). The carotenoids synthesised by *G. rubropertincta* are revealed through confocal microscopy to be stored completely in the cytoplasm and are not bound to the cell membrane. The carotenoids’ location was further confirmed by TEM. The phospholipid bilayer present on the exterior of the bacterial cell, genetic material that can be seen as a darkly stained body inside the cell, and the darkly stained cytoplasm could be seen through TEM. The cytoplasm getting stained in black indicates the presence of carotenoids in the cytoplasm of the cell. Being present in the cytoplasm makes it easier to bring the carotenoids out of the cell and separate it from other cell debris compared to those that are membrane bound. Therefore, extracting the carotenoids and purifying them should be easier when *G. rubropertincta* is used as the host organism.

## 5.0 CONCLUSION

To summarize, *Gordonia rubropertincta* is a gram-positive, actinobacteria that synthesises a broad range of carotenoids through various pathways. The carotenoids it synthesises are auto-fluorescent and present in the cell cytoplasm, so are easy to extract from the cell. It is a store house of different carotenoids. Each carotenoid can be used for different applications, and bacteria are easy to handle, study and maintain at lab to industrial scale compared to higher organisms like fungi, algae, and yeasts. Hence, this bacterium can be used to commercially produce carotenoids. The diversity of carotenoids it synthesises makes it an ideal organism to study carotenoids, their pathway(s), properties, applications, to metabolically engineer and increase or decrease production of a particular carotenoid, and channelise carotenoid synthesis into one or more particular pathway(s). This organism can be of high importance in industries to produce carotenoids commercially.

## ACKNOWLEDGEMENTS

The authors are thankful to The Director, CSIR-Indian Institute of Chemical Technology (IICT) for providing the facilities required for carrying out this research work (**IICT/Pubs./2023/097**). The authors are grateful to the Academy of Scientific and Innovative Research (AcSIR) for their immense support during this work. Also, sincere thanks to the Indian Council of Medical Research (ICMR) for providing Ms. Gayathri Vemparala with research fellowship (JRF-2019, HRD-22). Thanks to Dr. Mohana Krishna Reddy Mudiam, Dr. Rajkumar Banerjee, Narendra Kumar Nagendla for their help in analytical techniques. Thank you Dr. M. Srinivasa Rao and Dr. Gurrala Sheelu for reviewing and enhancing the article.

