## Supplementary Information for "‘Caroten-omics’ of pigments synthesised by *Gordonia rubropertincta* through High Resolution Mass Spectrometry and *in-silico* programming approach"

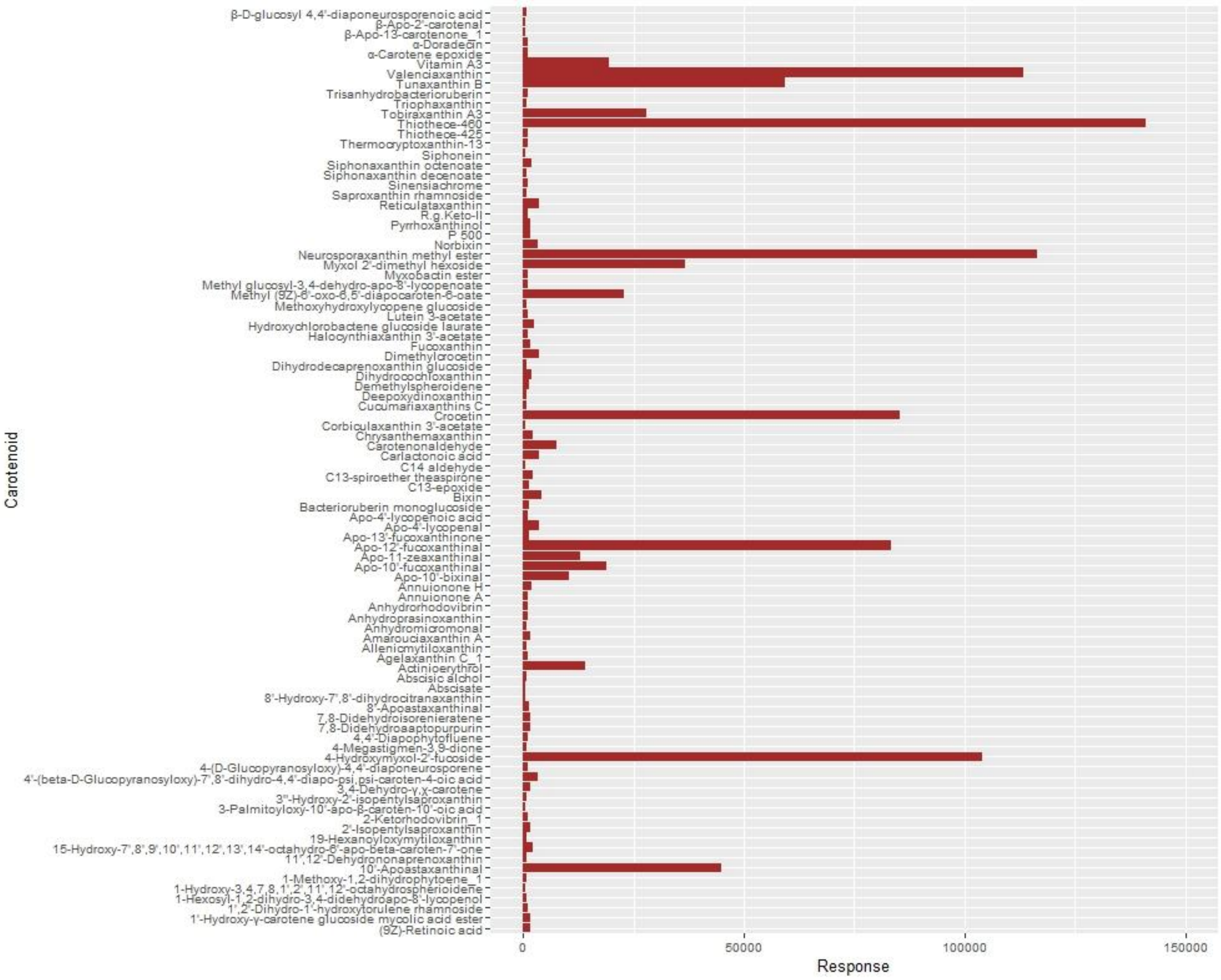


**Figure S1. The 93 carotenoids synthesized by *Gordonia rubropertincta* and the abundance of each carotenoid**

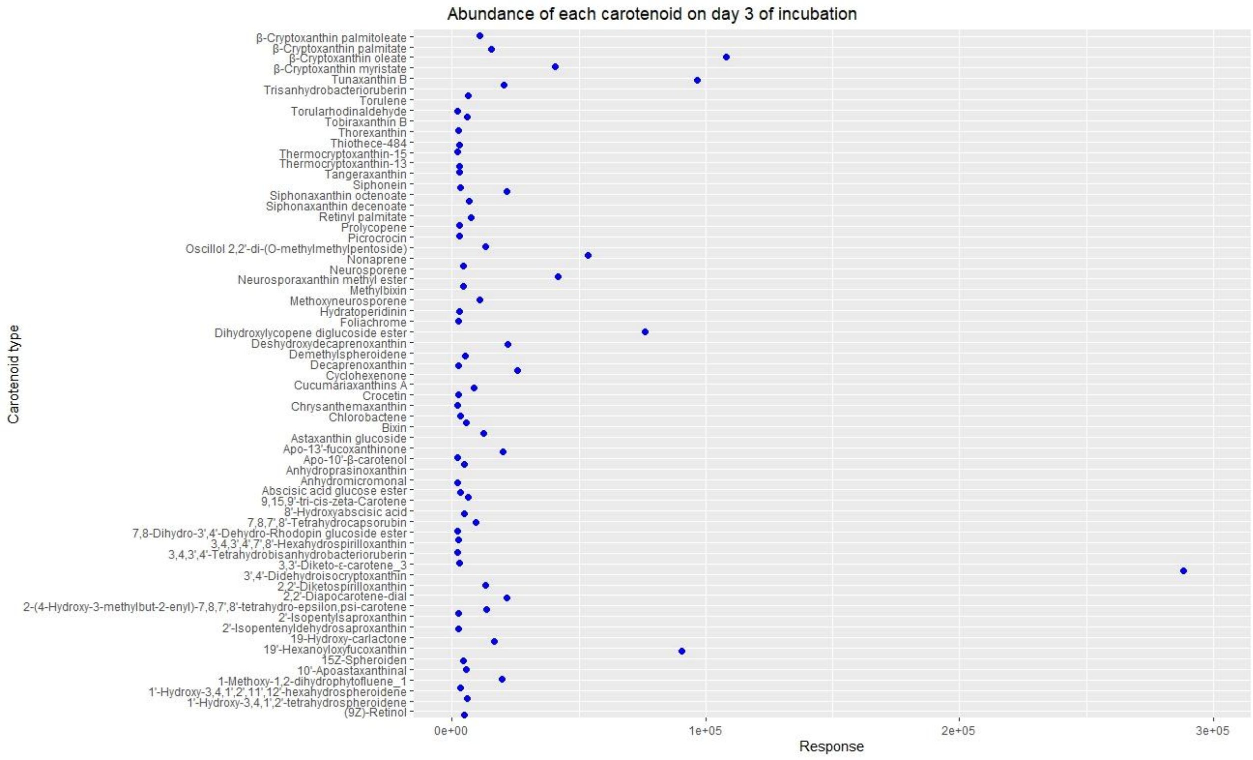


**Figure S2(a). The carotenoids synthesized by *Gordonia rubropertincta* on 3 days of incubation and the abundance of each carotenoid**

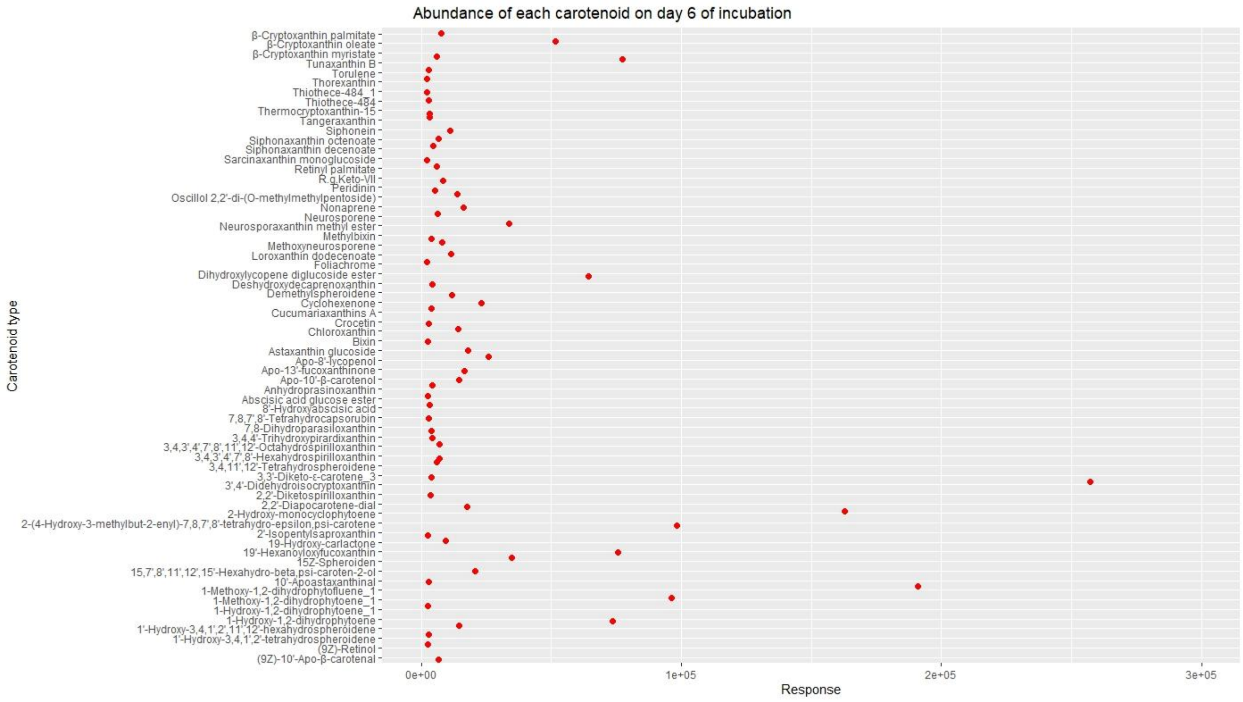


**Figure S2(b). The carotenoids synthesized by *Gordonia rubropertincta* on 6 days of incubation and the abundance of each carotenoid**

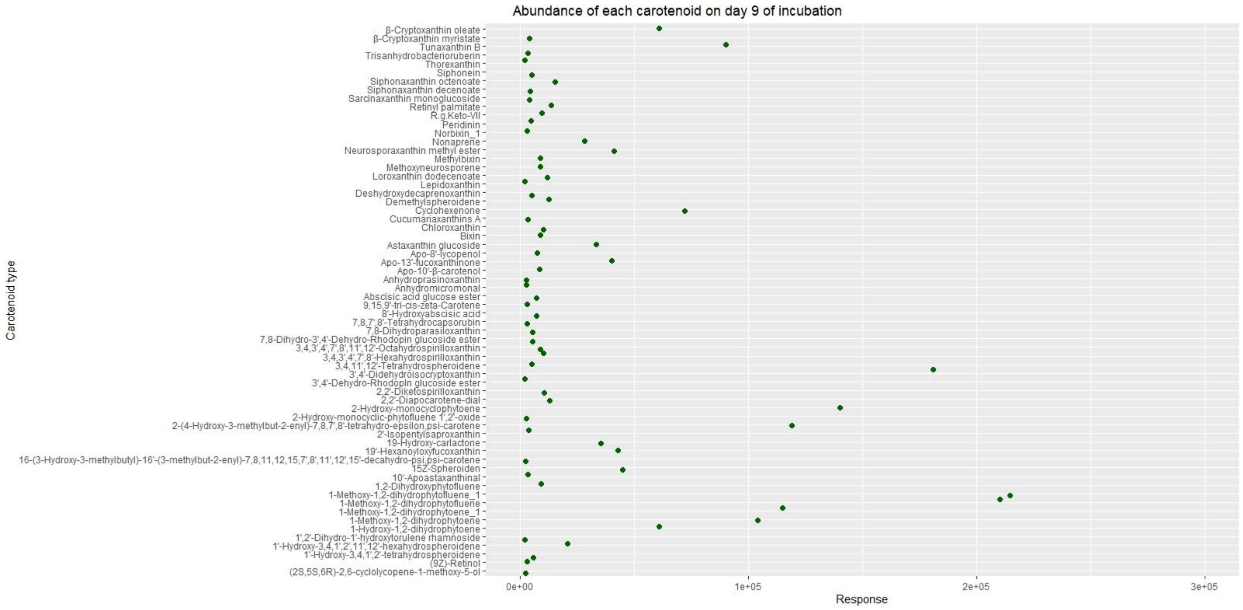


**Figure S2(c). The carotenoids synthesized by *Gordonia rubropertincta* on 9 days of incubation and the abundance of each carotenoid**

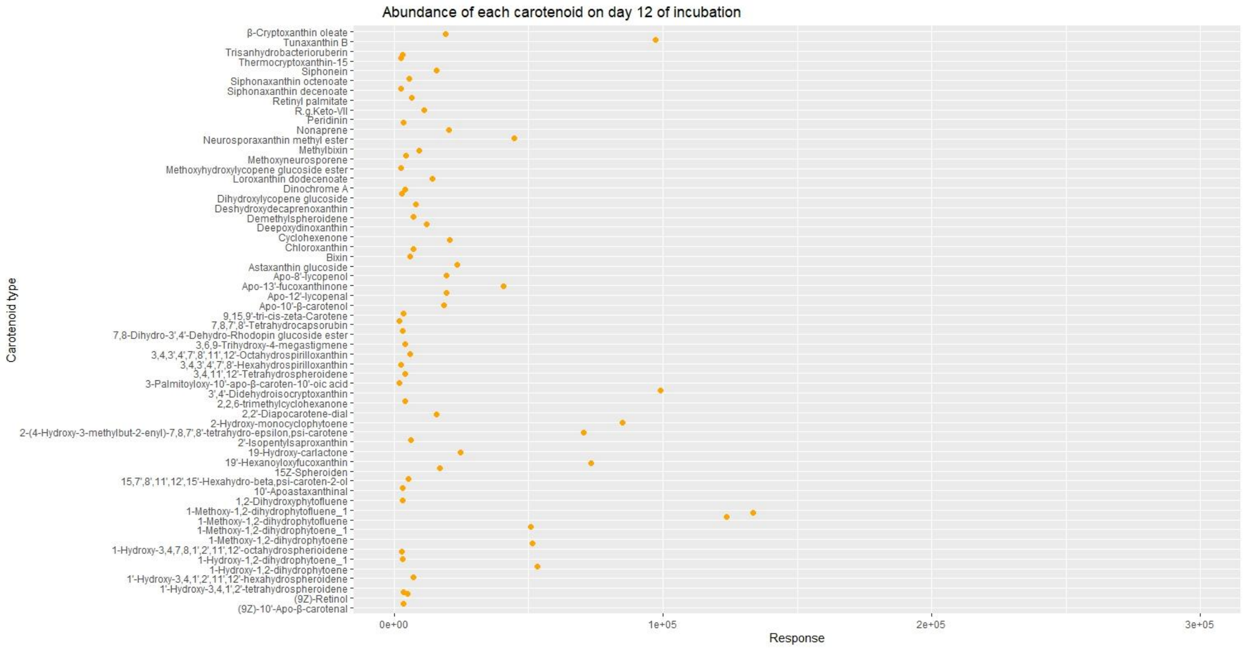


**Figure S2(d). The carotenoids synthesized by *Gordonia rubropertincta* on 12 days of incubation and the abundance of each carotenoid**

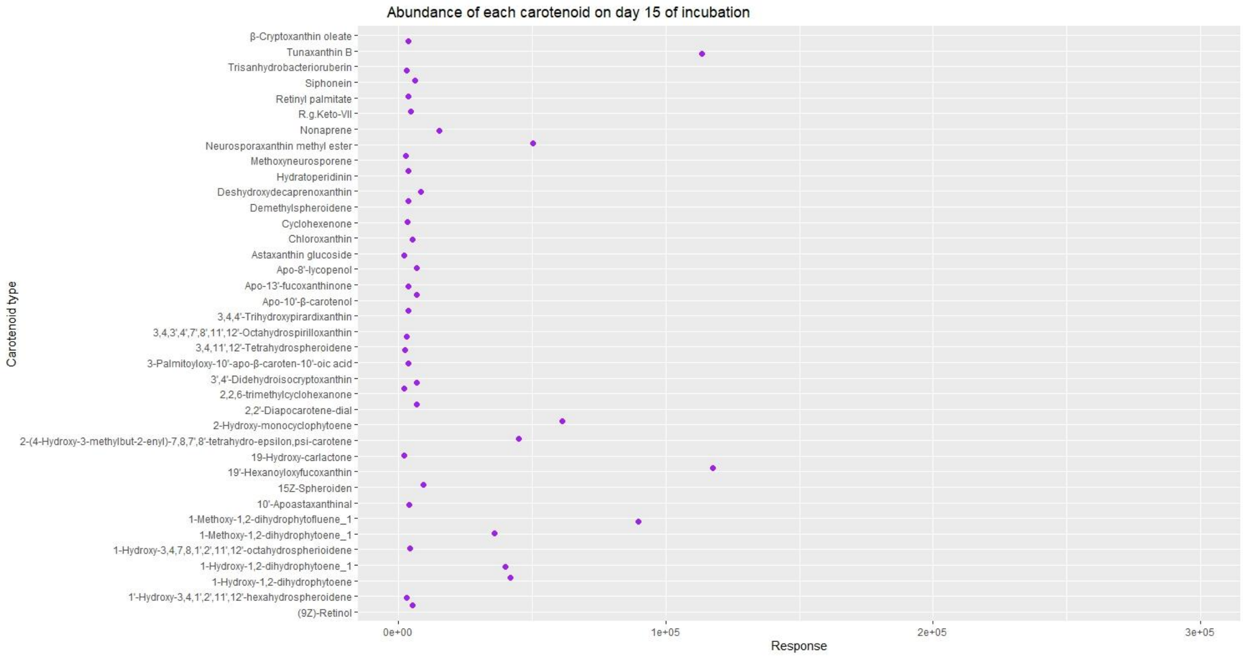


**Figure S2(e). The carotenoids synthesized by *Gordonia rubropertincta* on 15 days of incubation and the abundance of each carotenoid**
